# The transcriptional landscape of metastatic hormone-naïve prostate cancer

**DOI:** 10.1101/2024.05.20.594913

**Authors:** Natalia Martin-Martin, Saioa Garcia-Longarte, Jon Corres-Mendizabal, Uxue Lazcano, Ianire Astobiza, Laura Bozal-Basterra, Nicolas Herranz, Hielke van Splunder, Onintza Carlevaris, Mikel Pujana-Vaquerizo, María Teresa Blasco, Ana M. Aransay, Antonio Rosino, Julian Tudela, Daniel Jimenez, Alberto Martinez, Andrei Salca, Aida Santos-Martín, Sofía Rey, Aitziber Ugalde-Olano, David Gonzalo, Mariona Graupera, Roger R. Gomis, Joaquin Mateo, Miguel Unda, Enrique Gonzalez-Billalabeitia, Ana Loizaga-Iriarte, Isabel Mendizabal, Arkaitz Carracedo

## Abstract

Metastatic hormone-naïve prostate cancer (mHNPC) is an infrequent form of this tumour type that is characterized by metastasis at the time of diagnosis and accounts for 50% of prostate cancer-related deaths. Despite the extensive characterization of localized and metastatic castration resistant prostate cancer (mCRPC), the molecular characteristics of mHNPC remain largely unexplored. Here we provide the first extensive transcriptomics characterization of mHNPC. We generated discovery and validation bulk and single-cell RNA-Seq datasets and performed integrative computational analysis in combination with experimental studies. Our results provide unprecedented evidence of the distinctive transcriptional profile of mHNPC and identify stroma remodelling as a predominant feature of these tumours. Importantly, we discover a central role for the transcription factor SOX11 in triggering a heterotypic communication that is associated to the acquisition of metastatic properties. Our study will constitute an invaluable resource for a profound understanding of mHNPC that can influence patient management.

## Introduction

Prostate cancer (PC) is the most prevalent cancer type in men, and it is predominantly diagnosed in a localized stage, when curative-intended therapies are highly efficacious. Recurrent PC is subject to different means of androgen signalling inhibition and chemotherapy. Failure of these therapeutic approaches might result in an ultimately metastatic form of the disease (metastatic castration resistant PC, or mCRPC). The molecular landscape and driver alterations of localized PC and mCRPC have been extensively characterized^1–3^, contributing to build our current portrait of the disease. However, a minority of PC cases (estimated 10%)^4^ are diagnosed in a disseminated stage, offering a unique biological scenario where metastatic disease emerges prior to systemic therapy (metastatic hormone-naïve PC, mHNPC).

Despite the low incidence of mHNPC, these patients account for up to 50% of mortality by PC^5^. Prostatectomy is not a standard of care procedure in mHNPC cases, which limits biological sample availability and hampers the molecular characterization of this rare form of PC. A recent comprehensive study has highlighted extensive genomic intra-tumoural heterogeneity in mHNPC primary tumours^4^. This and previous studies support that mHNPC primary tumours genetically resemble advanced mCRPC specimens rather than primary locoregional tumours^4,6,7^ or, alternatively, present an intermediate profile^8,9^. Alterations that are common to both mHNPC and mCRPC include inactivation of tumour suppressor genes *TP53*, *PTEN* and *RB1*. In contrast, aberrations in androgen receptor *(AR)* gene are infrequent in PC^4,6,7,10^ despite reduced AR activity in these tumours^11^. Altogether, genomic evidence suggests that the aggressive features of mHNPC primary tumours predate therapy exposure. Nevertheless, this rare subtype of the disease remains severely uncharacterized at the molecular level. Improved clinical management strategies in mHNPC demand the generation of molecular resources that can impact guidelines for patient stratification and treatment.

Here, we present the first comprehensive transcriptome-wide characterization of primary tumours obtained from mHNPC patients using bulk and single-cell RNA sequencing technologies. For appropriate biological comparison, we contrasted primary tumours from patients with localized disease to untreated mHNPC primary tumours using needle biopsy-derived tissue. To account for the extensive inter-patient molecular heterogeneity that characterizes PC primary lesions^1–3^, we assembled and profiled two retrospective patient cohorts encompassing 110 individuals using bulk RNA-Seq. Extensive computational characterization of bulk RNA-Seq was combined with single-cell transcriptomics analysis of 15 cases, leading to the identification of *SOX11* as a potential driver of stroma remodelling and cancer cell dissemination in mHNPC.

## Results

### The transcriptional landscape of metastatic hormone-naïve prostate cancer primary tumours is highly divergent

To undertake a profound molecular characterization of mHNPC, we generated a series of patient cohorts with either localized disease or metastatic dissemination at diagnosis leveraging from retrospective studies in different Spanish hospitals, prospective biopsy collection and detailed clinical annotation of publicly available resources (**Fig. 1A**). Taking advantage of the availability of prostate biological material from formalin-fixed and paraffin embedded (FFPE) diagnostic needle biopsies, we first performed bulk RNA sequencing in a discovery cohort comprising 47 localized (LPC) and 31 mHNPC specimens (see workflow in **Fig. 1A** and **Supplementary Table 1** for cohort characteristics). We hypothesized that comparing primary tumour biopsies from metastatic and non-metastatic disease without the influence of therapy-induced selection pressure would offer us unique insights into the molecular determinants of PC aggressiveness. Unsupervised hierarchical clustering and principal component analyses (**Fig. 1B** and **Supplementary Fig. 1A**) revealed highly distinct transcriptomic profiles in these two primary tumour types, with notable interindividual variation within mHNPC tumours. We identified a large number of genes with significant transcriptomic differences between disease clinical states (5,147 differentially expressed genes or DEGs, |log2(fold change >1.5)| and false discovery rate (FDR) <0.05, **Fig. 1C** and **Supplementary Table 2**) that were organized into hundreds of biological processes differentially enriched (**Supplementary Table 3**). These data unravelled a profound molecular difference between LPC and mHNPC. Histopathological differences could represent a confounding factor in this analysis. Since mHNPC exhibited higher Gleason score (or ISUP grade^12^) (**Fig. 1D**), we took advantage of four publicly available cohorts encompassing 780 patients with localized disease with annotated Gleason score (**Supplementary Table 4**). Top 50 DEGs failed to cluster the patients according to pathological grading, suggesting that the observed differential expression is not driven by this parameter (**Supplementary Fig. 1B-E**).

**Fig. 1.**
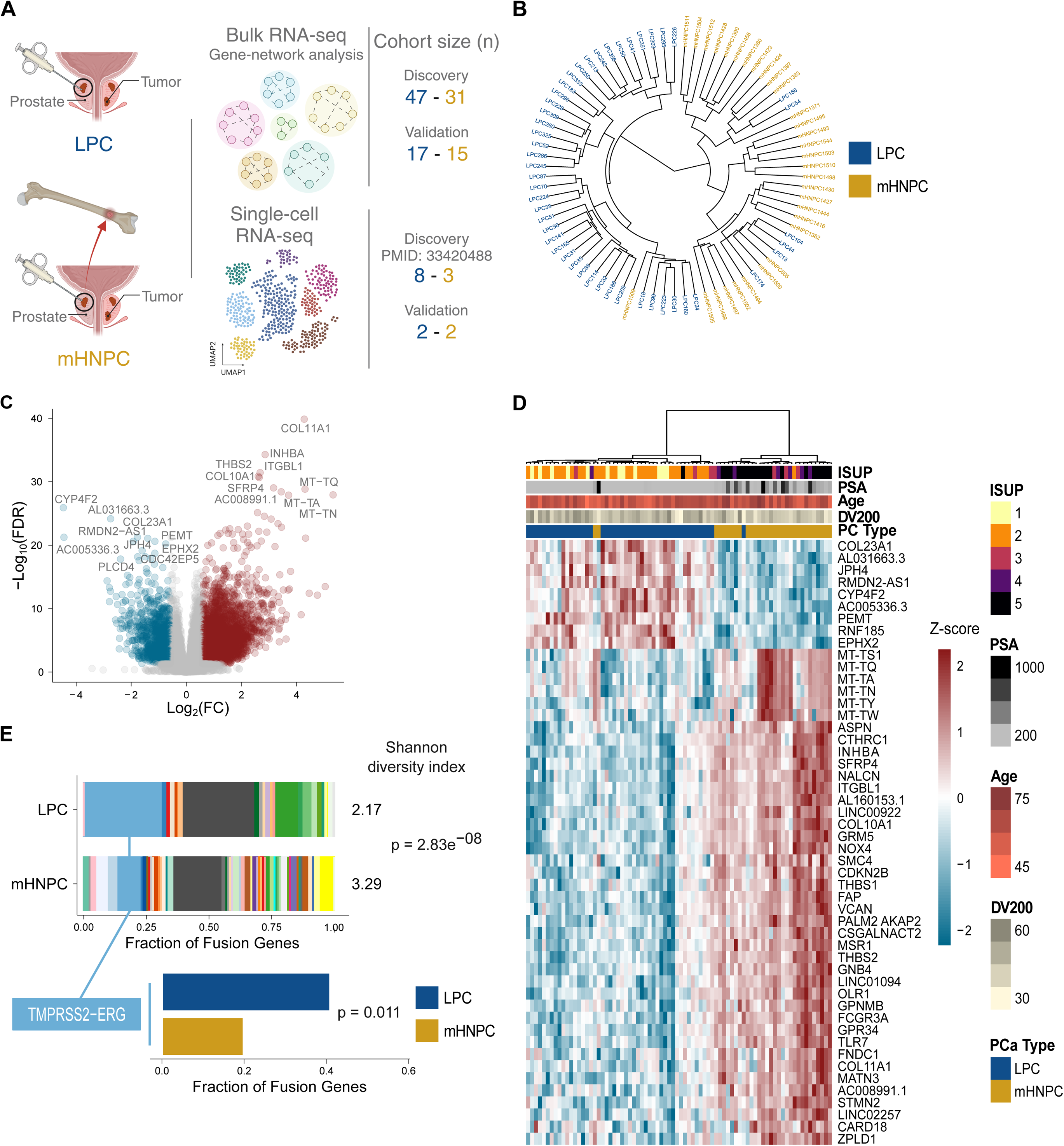
Divergent transcriptional profile of primary tumour specimens from localized and metastatic hormone-naïve prostate cancer patients. A. Overview of the cohorts of localized (LPC) and metastatic hormone-naïve (mHNPC) prostate cancer patients and transcriptomic datasets from primary tumours included in this study. **B.** Unsupervised hierarchical clustering of transcriptome-wide patterns of LPC and mHNPC primary tumours (discovery cohort). **C.** Volcano plot of differential gene expression analysis for the mHNPC vs LPC comparison. **D.** Heatmap and unsupervised clustering of the top 50 differentially expressed genes including technical (RNA quality score on FFPE samples, DV200) and clinical variables (age, PSA and ISUP). **E.** Gene fusion detection in LPC and mHNPC tumours from transcriptomic data. Diversity estimates according to Shannon diversity index are shown (p-value from Hutcheson’s t-test). The zoom shows TMPRSS2-ERG gene fusion class with the relative fractions in each tumour type (p-values from binomial test).

Transcriptional data can provide valuable information about genomic reorganization in these aggressive tumours^13^. In contrast to other cancer types, gene fusions represent the most common genetic alteration in PC, being present in over 70% of the cases^14,15^. We inferred gene fusions from the RNA-Seq data following the Trinity Cancer Transcriptome Analysis Toolkit (CTAT)^13^. In line with previous studies, >80% of patients in our cohort presented gene fusions (**Supplementary Table 5**). Despite similar total frequencies, the composition of fusion classes was markedly different in mHNPC, which exhibited remarkable diversity of fusion transcripts (Shannon Diversity Index, p-value = 2.83e^-08^, **Fig. 1E**). The increased diversity in genomic structural variation is in concordance with larger inter-patient variability observed in mHNPC transcriptome. Notably, while LPC showed a high frequency of *TMPRSS2-ERG* gene fusions, this event was underrepresented in mHNPC patients (19% in mHNPC vs 40% in LPC, binomial test, p-value=0.011, **Fig. 1E**). This translocation juxtapositions the androgen-regulated promoter of *TMPRSS2* with the coding region of the *ERG* oncogene. The existence of this gene fusion is associated to androgen-dependent tumours^16^, and the reduction in this event in mHNPC is consistent with decreased AR dependence and earlier development of resistance to androgen deprivation therapy^17^.

### Gene network analysis reveals increased stromal remodelling in mHNPC

Our transcriptomics results reveal profound gene expression differences between LPC and mHNPC, which represents an important challenge for the identification of driver candidate processes. Gene network analyses constitute powerful methods to identify coordinated activities of gene sets. To prioritize among the extensive list of potential biological processes underling the metastatic capacity of mHNPC primary tumours, we applied weighted gene correlation network analyses (WGCNA)^18^ (**Fig. 2A**, **Supplementary Fig. 2A-C**), where highly interconnected genes are clustered in modules. WGCNA identified 49 colour-coded modules, and 7 where strongly positively associated with the mHNPC phenotype (Pearson’s r > 0.6 and FDR<0.05, **Fig. 2B**). The “purple” module exhibited the highest coordinated overexpression in mHNPC (**Fig. 2B**, **Supplementary Fig. 2D** and **Supplementary Table 6**). This module (including 758 genes) incorporated the largest proportion of DEGs (77.7%), **Fig. 2B**) highlighting the convergence of gene-by-gene and network analyses on this gene set. Unsupervised hierarchical clustering revealed that the top 30 genes of the purple module robustly discriminated mHNPC from LPC (**Fig. 2C**). The DEGs showed larger intramodular connectivity than non-DEGs (Wilcoxon test, p-value=0.0013, **Fig. 2D**), implying that DEGs tend to be hub genes within the network. Interestingly, functional enrichment analysis identified processes related to stroma remodelling within the purple module in mHNPC (**Fig. 2E**).

**Fig. 2.**
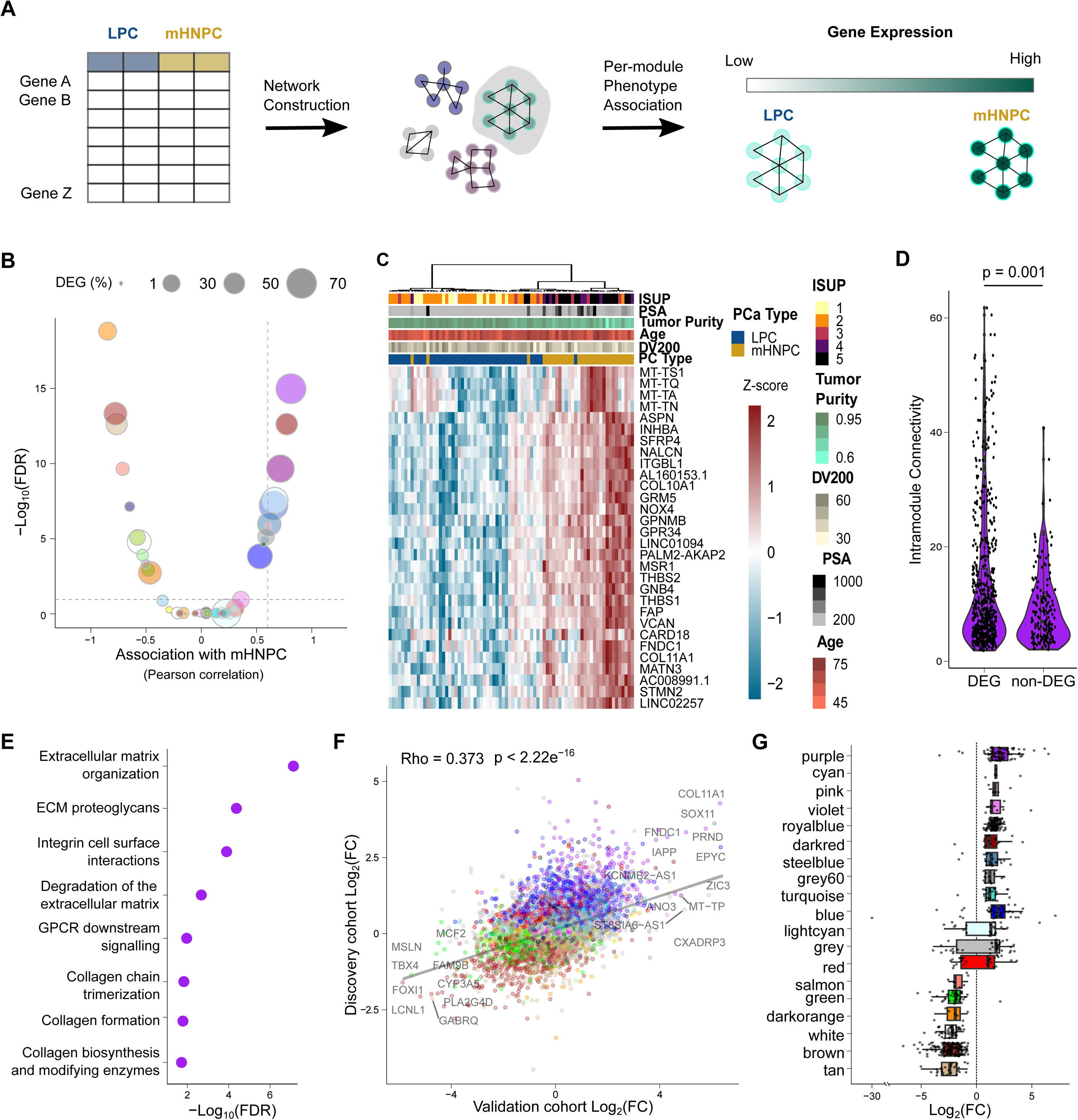
Gene-network analyses point at upregulation of stroma remodelling in mHNPC. **A.** Schematic representation of the gene module construction using weighted-gene network correlation analyses (WGCNA) and association with mHNPC and LPC phenotypes. **B.** Volcano plot showing the results of the association of each module with the mHNPC phenotype and the percentage of differentially expressed genes (DEGs) per module (dot size). Lines represent significance thresholds at Pearson’s r > 0.6 and FDR<0.05. **C.** Heatmap and unsupervised clustering of the top 30 DEGs in the purple module. **D.** Intramodular connectivity between the genes in the purple module (p-value from Wilcoxon test) according to differential expression (DEG status). **E.** REACTOME pathway enrichment results for the DEGs in the purple module. **F.** Correlation between the log_2_ fold changes of the differential expression analyses in the discovery (y-axis) and validation (x-axis) cohorts for all genes (p-values from Spearman’s correlation). **G.** Boxplot of the log_2_ fold changes in the validation dataset for DEGs grouped by the modules identified by WGCNA in the discovery dataset. Only modules with at least 5 genes are shown.

To validate our observations, we generated a validation patient cohort comprising primary tumour specimens from 17 LPC and 15 mHNPC patients from an independent hospital (**Supplementary Table 1**). In line with the results from the discovery cohort, transcriptome-wide variation robustly clustered the two disease types (**Supplementary Fig. 2E**, F). We identified 3,591 DEGs in mHNPC tumours in the validation cohort (|log2 (fold change >1.5)| and FDR <0.05, **Supplementary Fig. 2G**, **Supplementary Table 7**). Notably, the fold changes obtained in both cohorts were significantly correlated (**Fig. 2F**, Spearman’s rho=0.373, p-value<2.22e^-16^ for all genes and **Supplementary** Figure 2H, Spearman’s rho=0.833, p-value <2.22e^-16^ for DEGs in both datasets). When projecting WGCNA module ascription into the validation dataset, we corroborated that genes in the purple module were the most robustly upregulated in mHNPC (**Fig. 2F, G**). Our computational strategy allowed us to reduce the transcriptional changes observed in mHNPC into a subset of coordinated genes associated to stroma remodelling.

### Transcriptomics at single-cell resolution identifies mHNPC-specific heterotypic interaction programs coordinated by *SOX11*

Bulk transcriptomic analysis provides information of mRNA abundance with disregard to the cellular source, and obscure quantitative and functional cell type differences. To explore the compositional differences of the tumour biopsies, we employed *in silico* deconvolution methods that infer cell-type proportions from bulk transcriptomic data^19^. In the discovery cohort, the tumour purity estimates in mHNPC (namely, the relative abundance of the epithelial compartment) were significantly reduced, consistent with an increase in immune and stromal infiltration scores (**Supplementary Fig. 3A** and **Supplementary Table 8**). Analyses based on variance partitioning confirmed that the fraction of transcriptional variation explained by tumour purity was substantial and exhibited a larger contribution than the tumour type (mHNPC vs. LPC, **Supplementary Fig. 3B** and **Supplementary Table 8**). The validation cohort exhibited a similar trend towards a reduction in estimated tumour purity and increased non-immune stromal contribution, while the reduced statistical significance could be due to the smaller cohort size or to the differences in sample collection (RNA from the validation cohort was obtained from frozen tissue that was collected from needle biopsies for mHNPC and prostatectomies for LPC^20^) (**Supplementary Fig. 3C**, D). Changes in stroma remodelling could ascribed to qualitative (function) or quantitative (abundance) alterations, and the inference of qualitative stromal changes from bulk RNA-Seq pose a challenge^21^. To study functional tumour and stromal cell alterations in mHNPC, we analysed transcriptomic changes at single-cell resolution. We first leveraged a publicly available single-cell RNA-Seq cohort^22^ (**Fig. 1A** and **Supplementary Table 1**), and detected upregulated genes in mHNPC compared to LPC in 10 out of the 21 clusters identified (**Fig. 3A, B**, **Supplementary Fig. 4A**). The epithelial compartment, and specifically luminal cluster 6, exhibited the greatest number of DEGs (n=14, |log2(fold change >1.5)| and FDR<0.1, **Fig. 3B, Supplementary Table 9**). To confirm that cluster 6 contained tumour cells, we performed copy number inference^23,24^ (**Supplementary Fig. 4B**). Consistent with bulk transcriptomics analysis, stroma remodelling functions were overrepresented among the DEGs in cluster 6 (**Fig. 3C**). To test the coherence between single-cell and bulk differential expression analyses, we correlated the fold changes obtained for each strategy (minimum |log2(fold change>1.5|). Notably, genes in purple module showed the strongest correlation with cluster 6 in single-cell data and, conversely, genes within purple module where best represented in cluster 6 (**Fig. 3D, E** and **Supplementary Table 10**). Taken together, the integration of bulk and single-cell analyses identifies genuine molecular alterations in an epithelial tumour cell subset that is strongly associated to mHNPC.

**Fig. 3.**
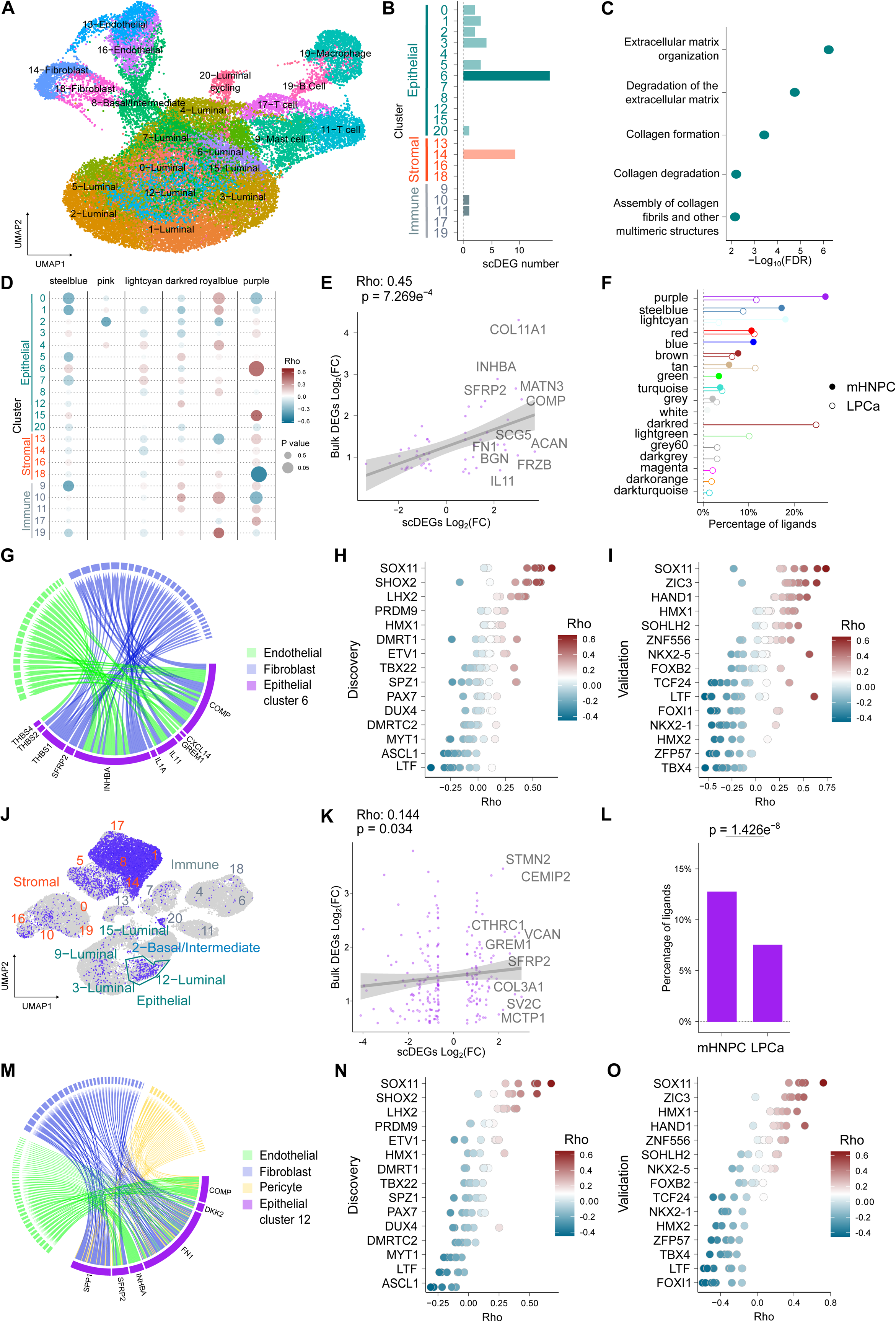
Single cell molecular deconstruction of HNPC. A. Visualization of single-cell discovery dataset (36,424 cells) using Uniform Manifold Approximation and Projection (UMAP). Colours code for the assignment of each cell to different clusters by graph-based clustering (Leiden algorithm). **B.** Number of differentially expressed genes (DEG) per single-cell cluster comparing mHNPC and LPC. **C.** Reactome pathway enrichment analysis for DEGs in cluster 6. **D.** Correlation analyses between the fold changes of differential gene expression (mHNPC vs LPC, minimum |log_2_(fold change)| >0.6) in bulk and single-cell for mHNPC– associated gene modules. Darkmagenta did not show any genes above the fold change criteria and is not represented in the figure. Dot colours show Spearman’s correlation values and dot size represents p-value. **E.** Correlation analysis of fold changes between bulk and single-cell data for genes in the purple module in cluster 6. **F.** Bar plot showing the module classification of the ligands identified in cell-cell communication between cluster 6 (sender) and non-immune stromal cells (receiver). **G.** Cord diagram of mHNPC-enriched ligands in the communication between cluster 6 and stromal cells that belong to purple module. Different stromal clusters were merged into categories for visualization. **H.** Spearman correlation of the purple ligands with transcription factors in bulk transcriptome data from mHNPC patients (discovery bulk cohort). **I.** Spearman correlation of the purple ligands with transcription factors in bulk transcriptome data from mHNPC patients (validation bulk cohort). **J.** Projection of 28 genes associated with cluster 6 in discovery dataset (i.e. markers that distinguish cluster 6 from other epithelial cells) in the validation cohort, exhibiting highest similarity with cluster 12 within the epithelial compartment. **K.** Correlation analysis of differential expression (fold change of mHNPC vs. LPC) between bulk and single-cell RNA-Seq in cluster 12 for purple module genes. **L.** Bar plot showing the results of cell-cell communication between cluster 12 (sender) and non-immune stromal cells (receiver) in the single-cell validation cohort for ligands with purple module membership . **M.** Cord diagram of mHNPC-enriched ligands in the communication between cluster 12 and stroma that belong to the purple module. Different stromal clusters were merged into categories for visualization. **N.** Spearman correlation of the purple ligands identified in the single-cell validation cohort with transcription factors in bulk transcriptome data from mHNPC patients (discovery bulk dataset). **O.** Spearman correlation of the purple ligands identified in the single-cell validation cohort with transcription factors in bulk transcriptome data from mHNPC patients (validation bulk dataset).

The presence of scDEGs in stromal clusters and the enrichment of mHNPC in stroma-regulatory processes suggested that molecular alterations in these tumour cells could govern stromal remodelling. We modelled intercellular communication between cluster 6 (as sender cells expressing ligands) and stromal cells (fibroblast and endothelial clusters as receiver cells expressing receptors), using NicheNet^25^. Among all processes showing differential connectivity between the two pathological conditions (LPC and mHNPC), we found an overrepresentation of ligands assigned to purple module (91/824, 11% of interactions in LPC, and 93/356, 26.12% in mHNPC, p-value = 1.004e^-10^, chi-squared test, **Fig. 3F**). Interestingly, some of the most differentially expressed genes replicated in bulk and single-cell cohorts were among the top ligands involved in the heterotypic communication process (**Fig. 3G** and **Supplementary Table 11**), including *SFRP2*, *INHBA*, *COMP* and *IL11*.

The transcriptional regulation of this ligand set belonging to purple module prompted us to search for transcriptional regulators that were perturbed in mHNPC and coordinatedly altered the expression of these genes. To this end, we designed a strategy based on gene-to-gene correlation analysis, establishing that ligand expression would correlate with the abundance of upstream transcription factors. To avoid the sparsity of single-cell data at single-gene level, we used the discovery bulk transcriptomic dataset and explored the correlation between all annotated transcription factors^26^ and the shortlisted purple module ligands. Interestingly, the SRY-Box Transcription Factor 11, *SOX11,* exhibited the strongest correlation with the ligand set (**Fig. 3H, Supplementary Table 12**). Of note, the corresponding cellular communication analyses between cluster 6 and the whole stroma (including immune clusters), pointed to a similar set of ligands (10/11 shared) and *SOX11* (**Supplementary Fig. 4C-D**). Importantly, this strategy was replicated using the bulk validation cohort (**Fig. 3I** and **Supplementary** Figure 4E).

We generated a validation single-cell RNA-Seq dataset from fresh needle biopsies from 2 LPC and 2 mHNPC patients (**Supplementary Table 1)**. In total, we profiled 36,424 cells and identified 21 clusters corresponding to epithelial, stromal and immune compartments (**Supplementary Fig. 4F**, G). In this new dataset, cluster 12 exhibited the greatest similarity to the cluster labelled as number 6 in the discovery single-cell dataset: i) luminal nature and shared cluster markers enriched in stroma remodelling, ii) large-scale chromosomal copy number alterations, and iii) significant association with purple WGCNA module regarding differential mHNPC expression (**Fig. 3J-K**, **Supplementary Fig. 4H-J)**. NicheNet-based cell-cell communication analysis between this cluster and the stromal compartment corroborated the over-representation of ligands belonging to the purple module among the mHNPC-enriched molecular communication processes (178/2349, 7.5% in LPC and 256/2001, 12.8% in mHNPC, p-value = 1.426e^-08^, chi-squared test, **Fig. 3L**). Some of the top ligands involved in this communication were concordant with the single-cell discovery dataset (*COMP, INHBA and SFRP2*), and new ones were also detected such as *FN1* and *SPP1* (**Fig. 3M** and **Supplementary Table 11**). Despite the expected variability in gene detection in highly sparse single-cell datasets, *SOX11* was corroborated as the transcription factor with the strongest correlation with this second set of ligands in mHNPC transcriptomes (**Fig. 3N, O**, discovery and validation bulk datasets, respectively, and **Supplementary Table 12**). In line with the discovery single-cell dataset, the ligand analysis including the immune stroma also pointed at *SOX11* (**Supplementary Fig. 4K-M**). Collectively, by integrating bulk and single-cell datasets from four independent cohorts, we consistently identified a tumour cell-intrinsic heterotypic communication program that is distinctive of mHNPC and potentially regulated by *SOX11*.

### *SOX11* promotes metastatic dissemination in PC

As a proof-of-concept of the potential relevant of the mHNPC transcriptional resource, we interrogated the relevance of SOX11 for the metastatic phenotype in PC. SOX11 belongs to the SOX transcription factor family. It plays crucial roles in stem cell function and tissue specification during embryogenesis but is largely absent in most differentiated adult tissues^27^. *SOX11* dysregulation is associated with oncogenesis, tumour progression, metastasis and therapy resistance^28–30^. Our integrative transcriptomic analyses unveiled that *SOX11* ranks prominently among the most overexpressed genes (**Fig. 2F, G**) and, importantly, it represents the top upregulated transcription factor (**Fig. 4A**). The observed elevation in *SOX11* levels and activity in mHNPC prompted us to study the contribution of this transcription factor to the metastatic phenotype. To this end, we engineered DU145 PC cells (which do not exhibit detectable levels of the transcription factor) to overexpress SOX11 (**Fig. 4B**, **Supplementary Fig. 5A**). Importantly, ectopic SOX11 expression resulted in increased mRNA and protein abundance of several purple module ligands identified in our scRNA-Seq analyses as contributors to the heterotypic communication between cancer cells and the stroma (**Fig 4C, Supplementary Fig. 5B** and **Supplementary Table 13**). Next, we inoculated control (3HA) or SOX11-overexpressing DU145 cells expressing orthotopically in the ventral prostate lobe of immunocompromised nude mice. Of note, both cell lines expressed GFP and luciferase for *in vivo* bioluminescence monitoring. Remarkably, SOX11-overexpressing PC cells exhibited increased metastatic capacity, with higher colonization of lumbar lymph nodes and bones (**Fig. 4D-E, Supplementary** Figure 5C). Of interest, the effects of SOX11 overexpression on cell number *in vitro* was negligible in contrast with the effect on tumour mass *in vivo* (**Supplementary Fig. 5D**, E). These data are consistent with the proposed role for this transcription factor in the activation of stroma remodelling processes, which would result in the manifestation of the phenotype in the presence of stroma but not in isolation (*in vitro* cancer cell culture).

**Fig 4.**
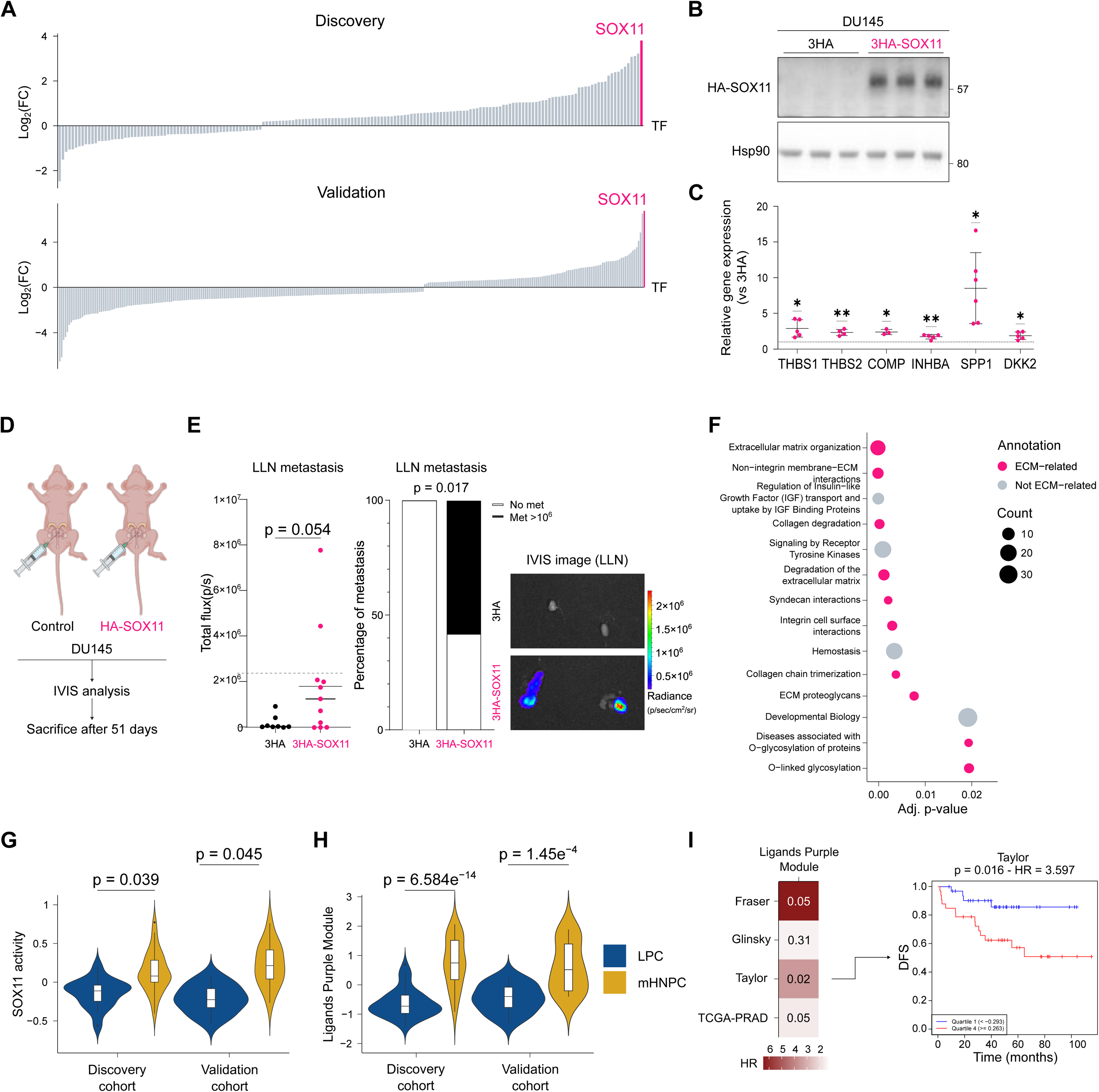
Proof-of-concept study of *SOX11* as a metastatic driver in prostate cancer. **A**. Waterfall plot representing the differential expression of transcription factors in mHNPC vs. LPC bulk RNA-Seq in discovery and validation cohorts. The position of *SOX11* is highlighted. **B.** Representative western blot in DU145 cells showing ectotopic HA-SOX11 expression (n=3). **C.** Evaluation of mRNA abundance in the indicated genes by quantitative real-time PCR upon ectopic HA-SOX11 expression (n=3-6). One sample t-test. **D**. Experimental design of the orthotopic injection of DU145 cells transduced with control and HA-SOX11-overexpressing lentiviral vectors into nude mice (n=5 for control, n=6 for 3HA-SOX11). **E.** Left, dot plot of lumbar lymph node (LLN) metastatic burden (day 51). One-tailed, Mann Whitney test. Middle, stacked bar plot of LLN metastasis appearance. One-tailed, Fisher test. Right, representative bioluminescence images of LLN metastasis. **F.** REACTOME pathway enrichment analyses of the genes upregulated upon *SOX11* overexpression (bulk RNA-Seq). **G.** Violin plots showing the levels of a SOX11-activity signature in the discovery and validation bulk RNA-Seq datasets. P-values from Wilcoxon test (two-tailed test for discovery and one-tailed for validation datasets, respectively). **H**. Violin plots showing the levels of a signature composed of genes encoding for purple module ligands identified in single cell RNA-Seq and correlated with SOX11 in the discovery and validation bulk RNA-Seq datasets. P-values from Wilcoxon test (two-tailed test for discovery and one-tailed for validation datasets, respectively). **I.** Hazard ratio (HR) and p-value of a signature based on the ligands used in (H) when analysing as endpoint biochemical recurrence in the indicated datasets and comparing patients within quartile 1 and 4. A representative Kaplan-Meyer curve is shown.

Taking advantage of this cellular model, we characterized transcriptomics alterations that emanate from *SOX11* overexpression by RNA-Seq. Functional enrichment analysis highlighted the regulation of stroma remodelling processes in *SOX11* overexpressing cells, consistent with our data in mHNPC (**Fig. 4F, Supplementary Fig. 5F, Supplementary Table 14**). Next, we generated a transcriptional signature of SOX11 activity, composed of 55 genes positively regulated by the transcription factor (|log2(fold change >1.5)| and FDR <0.05). This gene set was strongly associated to mHNPC in both the discovery and validation patient cohorts, confirming increased SOX11 activity in this aggressive form of PC (**Fig. 4G**). Our comprehensive transcriptional portrait of mHNPC reveals an unprecedented role for SOX11 in PC dissemination and supports its role in the acquisition of metastatic features in this disease. To ascertain the existence of elevated SOX11-dependent activity in association with other forms of aggressive PC, we took advantage of public transcriptomics dataset with extensive clinical annotation and follow up^3,15,31–34^. We did not observe consistent association of *SOX11* expression (*SOX11* mRNA abundance) or SOX11 inferred activity (the aforementioned 55-gene signature) with biochemical recurrence-free survival in patients diagnosed with localized PC (**Supplementary Fig. 5G-H**). Our data suggest that *SOX11* presents a genuine role ascribed to the pathogenesis of mHNPC. We hypothesized that despite the differential contribution of *SOX11* to LPC and mHNPC, the ligands identified in mHNPC could be contributing to the acquisition of aggressiveness features in localized disease governed by other transcriptional processes. To test this notion, we monitored the expression of the ligands reported to contribute to the heterotypic cell-cell communication in mHNPC under the control of SOX11. A gene expression signature based on the 13 ligands identified robustly discriminated mHNPC from LPC (**Fig. 4H**). Strikingly, this signature exhibited robust and consistent prognostic potential in patients with localized prostate cancer. This predictive capacity for biochemical recurrence after surgery is indicative of the presence of disseminated tumour cells with metastatic potential (**Fig. 4I**, **Supplementary Fig. 5I**).

## Discussion

Prostate cancer (PC) research has been dominated by the analysis of the most prevalent form of the disease, which refers to localized, prostate-confined, tumours (LPC). The scientific community has generated extensive clinical, biological and molecular information about PC based on the study of LPC^1–3^. Similarly, the alterations and therapeutic vulnerabilities of metastatic PC have been established upon the study of heavily treated PC patients that ultimately develop metastasis. Whereas this strategy has been tremendously valuable in the establishment of clinical guidelines in this tumour type, the translation of these evidence to mHNPC is uncertain. Considering that mHNPC cases, albeit infrequent, represent a large fraction of the mortality by PC^4,5^, it is of the utmost importance to characterize the molecular features of this form of prostate tumour to establish therapeutic strategies tailored to this aggressive disease.

Bulk transcriptomics offers valuable insight about predominant differences among biological conditions. However, complex biological specimens can pose a challenge due to the influence of mRNAs coming from different cell types^19^. In addition, when comparing profoundly different tumours, differential expression can prove insufficient to identify relevant molecular processes. Our implementation of WGCA^18^ combined with scRNA-Seq, together with the integrated use of bulk and single-cell information has enabled us to identify consistent stroma remodelling processes that can be illustrated by a set of SOX11-regulated ligands. On the one hand, the capacity of SOX11 to promote a metastatic phenotype suggests that the upregulation of this transcription factor contributes to the aggressiveness of mHNPC. On the other hand, the lack of upregulation of *SOX11* in LPC with varying degree of aggressiveness, and the capacity of the ligands identified in mHNPC to inform about the risk of biochemical recurrence suggests that the role of SOX11 in mHNPC might be exerted by other transcription factors in aggressive LPC.

In sum, this study provides the most comprehensive transcriptional portrait of mHNPC to date. The data generated and the various cohorts available will represent an invaluable resource to boost the molecular and biological deconstruction of mHNPC. We highlight the biological and molecular uniqueness of this aggressive form of PC and present proof-of-concept evidence of the value of this resource through the study of SOX11 as a transcription factor selectively activated in mHNPC that supports the acquisition of metastatic features.

## Methods

### Sample and patient disposition

Detailed information about the clinical and pathological characteristics of the cohorts included in this study are shown in Supplementary Table 1. Samples were labelled as LPC and mHNPC groups according to the patient metastatic status at the time of first diagnosis (M0, no distant metastasis present; M1, distant metastasis present, according to AJCC TNM stating system), upon review of electronic patient records, and based on standard of care imaging and clinical evaluations. The samples included in the discovery cohort for bulk RNA-Seq profiling (formalin fixed paraffin embedded specimens from primary tumour biopsies) were collected from patients at Basurto University Hospital in Bilbao (Spain). Sample collection was coordinated by the Basque Biobank. Samples with at least 5 (out of 12) positive needle biopsies were selected to ensure sufficient tumoural material for analysis. Tumour-rich regions were selected for RNA extraction by the pathologist (A.U-O). The ethics approvals for this project are CEIC-E 14-14 and 19-20. The patient cohort for bulk RNA-Seq for validation purposes was generated at the Morales Meseguer University Hospital in Murcia (Spain). Fresh frozen OCT-embedded tissue specimens were obtained from prostatectomy for LPC patients and from needle biopsies for mHNPC patients. The ethics approval for this project was CEIC-HMM 1/18. The single-cell validation cohort was generated from patients recruited in Basurto University Hospital in Bilbao (LPC patients) and Vall d’Hebron University Hospital in Barcelona (mHNPC patients), under IRB-approved protocols CEIC-E 14-14 and 19-20, and PRAG5248 respectively.

### Bulk RNA-Seq analyses on discovery cohort

We performed RNA-Seq (paired-end 150bp reads, 86M reads per sample in average) from formalin-fixed paraffin-embedded (FFPE) specimens derived from primary tumours in patients presenting localized and metastatic prostate cancer at the time of diagnosis (recruited at the Basurto University Hospital, Spain). Sequencing libraries were prepared using SMARTer Stranded Total RNA-seq Kit v2 – Pico Input Mammalian kit (Takara Bio USA, Cat.# 634411) and following the user manual (Rev. 063017). Based on the library specifications and read length of 150bp, reads were specifically trimmed for 3 specific nucleotides using cutadapt v3.5 sotfware as follows. First, we removed the first 3 nucleotides of R2 (cutadapt -U 3 --pair-filter=any --minimum-length=30), then removed adapters from both reads using TruSeq adapters (cutadapt -b file:cutadapt_TruSeq_CD_R1.fa -B file:cutadapt_TruSeq_CD_R2.fa --minimum-length=30 --pair-filter=any) and finally remove the last 3 nucleotides in R1 (cutadapt -u -3 -q 10 --pair-filter=any --minimum-length=30). After quality control using fastQC we mapped the reads to the human genome (Ensmbl GRCh38.v94) using STAR^35^ version=2.7.0e with parameters: --outFilterMultimapNmax 1 --twopassMode Basic.

We identified differentially expressed genes (DEGs) between the two tumour types using DESeq2 (V1.34.0)^36^ and applying variance stabilizing transformation (VST). We pre-filtered genes with less than 10 counts per smallest group size. We used DV200 (representing the percentage of RNA fragments larger than 200 nucleotides, as a measure of RNA quality in FFPE samples) and age as covariates in the analyses. Cell composition analyses were performed using ESTIMATE^19^ R package (v1.0.13). Variance partitioning was performed using variancePartition^37^ R package (v1.24.1). Enrichment analyses were performed using gprofiler2 (v0.2.1)^38^ . Gene fusions were detected with the Trinity CTAT Fusion workflow^13^ using docker trinityctat/starfusion (CTAT library version Apr062020). First, we ran STAR-Fusion to identify candidate fusion transcripts based on discordant read alignments. The predicted gene fusions were ’in silico validated’ using FusionInspector. WGCNA^18^ was run using R package “wgcna” (v1.64.1 in docker mochar/wgcna). We used VST counts as input and automatic, one-step method with the following parameters: corType=”bicor”, deepSplit=2, networkType = ”signed”.

### Bulk RNA-Seq data analyses on validation cohort

We performed RNA-Seq (paired-end 150bp reads, 46M reads per sample in average) from snap frozen OCT primary tumours collected from patients at the Morales Meseg uer University Hospital. Sequencing libraries were prepared using the TruSeq Stranded mRNA LibraryPrep kit (Illumina Inc. Cat. # 20020594) and TruSeq RNA CD Index Plate (Illumina Inc. Cat. # 20019792). Reads were trimmed for Illumina universal adapter using cutadapt: cutadapt -a “AGATCGGAAGAGCACACGTCTGAACTCCAGTCA” -A “AGATCGGAAGAGCGTCGTGTAGGGAAAGAGTGT” -j 0 -q 10 -m 30). Mapping and differential expression analyses were performed as described above for the discovery cohort, using RIN values in DESeq2 (instead of DV200) as covariates and applying no filtering for minimum reads in this validation cohort.

### Analyses on single-cell discovery cohort

We accessed single-cell RNA-Seq data (10X Genomics) from GEO with accession ID GSE141445. The data consisted of processed count matrix on 13 patients with filtered mitochondrial genes. We used DoubletFinder (version 2.0.3)^39^ to identify doublets. The samples were SCTransformed by v2 regularization by Seurat^40^ . To integrate the samples, we used 3,000 anchor genes using the Canonical Correlation Analysis reduction method. We performed the linear dimensional reduction and the Uniform Manifold Approximation and Projection (UMAP) analysis with the top 13 PCs. The clustering was performed using a resolution of 0.8. Cluster markers were identified using FindAllMarkers function and cell types were annotated based on well-stablished markers: T cells (*CD3D*, *CD3E*, *CD3G* and *PTPRC*), B cells (*CD79A*, *CD79B*, *IGKC* and *MS4A1*), macrophages (*CD14*, *CD68*, *CSF1R*, *FCGR3A* and *LYZ*), mast cells (*KIT*, *MS4A2*, *TPSAB1* and *TPSB2*), fibroblasts (*COL1A1*, *COL1A2*, *COL3A1*, *DCN*, *RGS5* and *ACTA2*), endothelial cells (*CDH5*, *ENG*, PECAM1 and *VWF*), epithelial cells (*AR*, *EPCAM*, *KRT5*, *KRT8*, *KRT14*) and cell cycling marker genes (*BIRC5* and *CENPF*).

Differential expression analysis per cluster was performed using a pseudo-bulk approach with DESeq2 (version 1.36.0) using only the singlets (Benjamini–Hochberg FDR < 0.1). UCell (version 2.0.1)^41^ was used for evaluating gene signature scores, based on the Mann-Whitney U statistic. CopyKAT^23^ R package (version 1.1.0) was used to identify genome-wide aneuploidy at 5MB resolution in single cells, non-epithelial clusters 9, 10, 11, 13, 14, 16, 17, 18, 19 were used as reference. Cell-cell communication analyses were performed using NicheNet^25^ R package (v1.1.1). For prioritization based on differential expression, we used Wilcoxon test on SCT counts (log fold change of 0.25 and expression pct of 0.1). The transcription factor list was obtained from https://humantfs.ccbr.utoronto.ca/download.php.

### Generation of the validation single-cell RNA-Seq dataset

The tumour was washed with cold phosphate-buffered saline (PBS; GIBCO) and minced into fragments under 0.4 mm. The tissue was chemically dissociated in a filter-sterilized with Liberase TH solution (2.5mg/ml; REF QZY-5401135001, Thermo Scientific) in complete media composed of Advanced DMEM/F-12 (REF 12-634-010, Fisher Scientific), Pen/strep 1% (REF 15-140122, GIBCO), MgCl2 (5mM, REF M4880, Sigma Aldrich), DNAse I (200 μg/ml, REF 11284932001, Sigma Aldrich) and Y-27632 dihydrochloride (10 μM; AbMole) at 37°C shaking at 800 r.p.m for 40 minutes. The tissue solution was manually disaggregated with a pipette every 10 min during the incubation followed by centrifugation at 2400xg for 5 min. Next, a second chemical dissociation was performed with TrypLE (REF 12605010, Thermo Scientific) supplemented with Y-27632 dihydrochloride under constant pipetting for 2 minutes. The inactivation of TrypLE was done by adding FBS (GIBCO) and centrifuged at 2400xg for 5 minutes. The cell pellet was resuspended in PBS, filtered through 40μm filter and washed with PBS. The suspension was centrifuged at 2400xg for 5 minutes and resuspended in 200ul of PBS. Cells were counted on a Neubauer chamber and 20000 cells were then centrifuged at 2400xg for 5 minutes and resuspended in PBS for scRNA-Seq analysis according to manufacturerś indications (targeted number of cells: 10000).

We use the 10X Chromium Controller to encapsulate and barcode single cells (Reagent Kits v3.1). The sequenced data were mapped to human reference genome GRCh38 with Cell Ranger (version 7.0.1). Seurat package (version 4.3.0) in R (version 4.2.1) was used for downstream analyses. Low quality cells expressing less than 200 unique genes, showing novelty score <80% (the ratio of number of genes over number of UMIs) or >20% of mitochondrial genes were excluded. Only genes expressed in more than 10 cells were shortlisted for subsequent analyses. We integrated and normalized data as explained above. For this dataset, during the normalization step we regressed out the mitochondrial percentage and we used a resolution of 0.6. Cell-type annotation was based on the following markers: T cells (*CD3D*, *CD3E*, *CD3G* and *PTPRC*), B cells (*CD79A*, *CD79B*, *IGKC* and *MS4A1*), macrophages (*CD14*, *CD68*, *CSF1R*, *FCGR3A* and *LYZ*), mast cells (*KIT*, *MS4A2*, *TPSAB1* and *TPSB2*), fibroblasts (*COL1A1*, *COL1A2*, *COL3A1*, *DCN*, *RGS5* and *ACTA2*), endothelial cells (*CDH5*, *ENG*, *PECAM1* and *VWF*), epithelial cells (*AR*, *EPCAM*, *KRT5*, *KRT8* and *KRT14*) and schwann cells (*S100B*, *NRXN1* and *SOX10*). CopyKAT R package (version 1.1.0) was used to identify genome-wide aneuploidy at 5MB resolution in single cells, non-epithelial clusters 0, 1, 4, 5, 6, 7, 8, 9, 10, 11, 13, 14, 16, 17, 18, 19, 20 were used as reference.

### Cell culture

The human prostate carcinoma cell lines used were purchased from Leibniz-Institut DSMZ (Deutsche SammLung von Mikroorganismen und Zellkulturen GmbH), which provided authentication certificate: DU145 (ACC261). Human embryonic kidney 293FT cells were generously provided by the laboratory of Dr. Rosa Barrio. DU145 and HEK 293FT were cultured in Dulbecco’s Modified Eagle Medium (DMEM, Gibco) supplemented with 10% (v/v) fetal bovine serum (FBS, Gibco) and 1% (v/v) penicillin-streptomycin (Gibco). Cells were maintained at 37°C and 5% CO2 in a humidified atmosphere. Possible mycoplasma contamination was routinely monitored across all cell lines using the MycoAlert detection Kit (Lonza; LT07-318).

### Generation of stable cell lines

For constitutive *SOX11* overexpression (3HA-SOX11) previously generated DU145 GFP-Luc cells were infected with a modified Lenti-EFS/P2A-blast plasmid donated by Dr. James D Sutherland (Addgene: #208041) in which the TURBO-ID region was substituted by a 3HA-tag and the murine form of *Sox11* (obtained from Addgene: #120387) was cloned after the tag (between restriction sites for EcoRI and BamHI). The same plasmid without the *SOX11* insert was used as a control (3HA-ctrl). The infection was performed using standard procedures: 293FT cells were transfected with the appropriate lentiviral vectors using the calcium phosphate method and the viral supernatant plus protamine sulfate were used to infect the DU145 cells. Cells were selected with blasticidin (10µg/ml) for 5 days (blasticidin was renewed after the first 3 days)^42^.

### Cellular and molecular assays

Proliferation assays were performed by plating 5000 DU145 cells in triplicate in 12-well dishes and fixing in formalin at the indicated time points^43^. Cells were stained with crystal violet as previously described and quantified after the resuspension of the crystals in 10% acetic acid by measuring the absorbance at 595nm. Protein extraction and Western blot were performed as previously described^44^. Briefly, samples were run in 4-12% gradient Nupage precast gels (Life Technologies, WG1403BX10) in MOPS buffer. Primary antibodies for HA (Cell Signalling Technologies, 3724), SOX11 (Cell Signalling Technologies, 58207) and HSP90 (Cell Signalling Technologies, 4874S) were used at a 1:1000 dilution. Secondary anti-rabbit antibody (Jackson ImmunoResearch, 111-035-144) was used at a 1:4000 dilution.

RNA was automatically extracted using the Maxwell RSC (Promega) platform and extraction kits (Promega, AS1390), following manufacturer’s instructions. Complementary DNA (cDNA) from 1000 ng of the extracted RNA was synthesized with Maxima™ H Minus cDNA Synthesis Master Mix (Invitrogen Ref: M1682). Synthesized cDNA was diluted 1:3 and QS5 or QS6 (Life Technologies) systems were used for performing RT-qPCR analyses. Gene expression is normalized to GAPDH expression. Applied biosystems TaqMan probes or primers with their corresponding Universal Probe Library (Roche) probes were used: m*SOX11* (Probe 11, F: ACAACGCCGAGATCTCCAAG, R: TGAACGGGATCTTCTCGCTG), *THBS1* (Hs00962908_m1), *THBS2* (Hs01568063_m1), *COMP* (Hs00164359_m1), *INHBA* (Hs01081598_m1), *DKK2* (Hs00205294_m1), *SPP1* (Hs00959010_m1), *FN1* (Probe 25, F: GGGAGAATAAGCTGTACCATCG, R: TCCATTACCAAGACACACACACT), *MEX3A* (Hs00863536_m1), *GAPDH* (Hs02758991g1).

### Prostate orthotopic xenograft model of metastasis

The procedures for animal experimentation were carried out in compliance with the ethical guidelines defined by the Biosafety and Animal Welfare Committee at CIC bioGUNE, following AAALAC recommendations. Mice were maintained in a controlled environment, with standard 12:12 light:dark cycles, 30-50% of humidity and controlled temperature at 22±2°C. Diet and water were provided *ad libitum*. At experimental endpoint, mice were sacrificed by CO2 inhalation followed by cervical dislocation.

Orthotopic models were generated by injection of either DU145 GFP-Luc 3HA-ctrl or DU145 GFP-Luc 3HA-SOX11 into the ventral lobe of the prostate of nude mice. 2x10^6^ cells were injected per mice in 50 µl of PBS:Matrigel (Corning, 356231) (70:30) mixture. Mice were imaged by IVIS right after the surgery to check that the injection was properly performed as well as to check for accidental dissemination (Day 0) and were subsequently monitored by IVIS on a weekly basis. Mice were sacrificed after 51 days and prostate cancer cell dissemination to distant organs was assessed by ex vivo monitoring of selected organs by IVIS.

### Bulk RNA-Seq data analyses on SOX11 overexpression

We performed RNA-Seq (paired-end 100bp reads, >43M reads per sample in average) from DU145 cells. Sequencing libraries were prepared using the TruSeq Stranded mRNA LibraryPrep kit (Illumina Inc. Cat. # 20020594) and TruSeq RNA CD Index Plate (Illumina Inc. Cat. # 20019792). Reads were trimmed for Illumina universal adapter using cutadapt: cutadapt -a “AGATCGGAAGAGCACACGTCTGAACTCCAGTCA” -A “AGATCGGAAGAGCGTCGTGTAGGGAAAGAGTGT” -j 0 -q 10 -m 30). Mapping and differential expression analyses were performed as described above, applying the filtering for minimum reads but setting independent filtering parameter from DESeq2 to false. Enrichment analyses were performed using g:Profiler. The signature for SOX11 activity (comprising 55 upregulated genes) was obtained by doing the average of the z-score values of these genes per sample.

### Proteomic analysis of SOX11 overexpression Sample preparation

Protein was extracted by incubating cells in a buffer containing 7M urea, 2M thiourea, and 4% CHAPS. Samples were incubated in this buffer for 30 min at RT under agitation and digested following the FASP protocol described by Wisniewski et al. 2009 with minor modifications (PMID: 19377485). Trypsin was added in 50mM ammonium bicarbonate to a trypsin:protein ratio of 1:10, and the mixture was incubated for overnight at 37°C. Peptides were dried out in an RVC2 25 speedvac concentrator (Christ) and resuspended in 0.1% FA. Peptides were desalted and resuspended in 0.1% FA using C18 stage tips (Millipore) prior to acquisition.

### Mass spectrometry analysis

The resulting peptides were loaded onto an EvoSep One (EvoSep) chromatograph coupled on-line to a TIMS ToF Pro mass spectrometer (Bruker), that uses Parallel Accumulation Serial Fragmentation (PASEF) acquisition to provide extremely high speed and sensitivity. 30 SPD protocol (approx. 44min. runs) was used, under default Evosep settings. Data-independent acquisition (DIA) was used for the acquisition of data.

### Protein identification and quantification

The obtained data was then processed with DIA-NN^45^ software for protein identification and quantification. Searches were carried out against a database consisting of human protein entries from Uniprot in library-free mode. Search parameters were 20ppm precursor and fragment tolerance, 0.05 carbamidomethylation of cysteines as fixed modification and oxidation of methionines as variable modification.

Quantitative protein data was loaded onto Perseus software (free software from Max Plank Institute, Munich)^46^. This program was used for the differential protein abundance analyses. For this purpose, protein abundance data was log2 transformed, filtered based on reproducibility (proteins present in at least 70% of the samples of one of the groups were kept in the analysis) and imputated (missing values were substituted by abundances randomly taken from the 10% least abundant proteins in each sample). A Student’s T-test was applied and proteins with a p<0.05 were considered as significantly differential.

### Statistical analysis on *in-vitro* and *in-vivo* experimental data

Sample size was not predetermined using any statistical method and experiments were not randomized. Investigators were not blinded during experiments or outcome assessment. All the experiments were performed with at least three biological replicates. N values represent the number of independent biological experiments, the number of individual mice or the number of patient samples.

For *in vitro* experiments one sample t-test was used to compare the values normalized to the control with a hypothetical value of 1 and results are presented as mean ± standard deviation. For *in vivo* experiments one-tail, Mann-Whitney test was used and results are presented either as the mean in a dop plot or as stacked bar plot where one-tail, Fisher-test analysis was applied in the contingency table. The confidence level used for all statistical analysis was 95% (p-value = 0.05). Outliers were determined an interval spanning over the mean plus/minus two standard deviations.

## Data and script availability

All data generated in this project (raw and processed) are available in GEO and SRA (data submitted, accession ID pending) including bulk, single-cell RNA-Seq and proteomics. Code for data analyses are publicly available in Github at https://github.com/imendizabalCIC/The-transcriptional-landscape-of-metastatic-hormone-naive-prostate-cancer-main.

## Acknowledgements

This study was predominantly funded by the Spanish Association Against Cancer (AECC, GCTRA18006CARR to Carracedo, Gomis, Unda, Graupera and González-Billalabeitia as PIs) and AstraZeneca Jóvenes Investigadores 2023 Award (To Carracedo, Mateo, Mendizabal and Herranz). The work of A. Carracedo is supported by the Basque Department of Industry, Tourism and Trade (Elkartek), the BBVA foundation (Becas Leonardo), the MICINN (PID2022-141553OB-I0 (FEDER/EU); Fundación Cris Contra el Cáncer (PR_EX_2021-22), Severo Ochoa Excellence Accreditation (CEX2021-001136-S), European Training Networks Project (H2020-MSCA-ITN-308 2016 721532), the, Fundación Jesús Serra, iDIFFER network of Excellence (RED2022-134792-T), and the European Research Council (Consolidator Grant 819242). CIBERONC was co-funded with FEDER funds and funded by ISCIII. M. Graupera is supported by Worldwide Cancer Research (WCR 21-0159). I. Mendizabal is supported by a CRIS Contra El Cancer Foundation to I. Mendizabal (PR_TPD_2020-19). J. Mateo is supported by CRIS Talent Award (TALENT20-10) and a Department of Defense CDMPR Physician-Science Award (PC220307). VHIO authors would like to acknowledge the Spanish State Agency for Research (Agencia Estatal de Investigación) for the financial support as a Center of Excellence Severo Ochoa (CEX2020-001024-S/AEI/10.13039/501100011033), the Cellex Foundation for providing research facilities and equipment, FERO Foundation and the CERCA Programme from Generalitat de Catalunya for their support. Laura Bozal was supported by the AECC Foundation (POSTD19048BOZA). H. van Splunder received funding from the European Union’s Horizon 2020 research and innovation programme under the Marie Skłodowska-Curie grant agreement No 955951. R. Gomis and M.T. Blasco were supported by the BBVA Foundation, Fundación Científica AECC (PRYGN223207GOMI), and MICINN (PID2022-143093OB-I00; FEDER/EU). We thank Monika Gonzalez, Laura Bárcena and Nuria Macias-Cámara from the Genome Platform Analyses at the CIC bioGUNE and Lidia López Jiménez from the single-cell Unit at the Josep Carreras Institute for generation of single-cell RNA-Seq libraries. We thank James D. Sutherland for the donation of the overexpression plasmid. We thank Felix Elortza and Mikel Azkargorta from CIC bioGUNE proteomics platform for proteomics sample processing and analysis. We thank Edurne Berra, Amaia Ercilla and Amaia Zabala for the revision of the manuscript. Illustrations were created with BioRender.

## Authorś contributions

S.G.-L., U.L. and I.M. performed bioinformatic analyses on bulk and single-cell transcriptomic datasets and contributed to the preparation of the figures for the manuscript. I.M. supervised the work of S.G.-L. and U.L. J.C.-M. and N.M.-M. performed most *in vitro* and *in vivo* assays, performed the data analysis and contributed to the preparation of the figures of the manuscript. N.M.-M. supervised the work of J.C.-M.I.A., M.P.-V., O.C., and T.B. provided technical support with the *in vivo* experiments. A.M.A. performed wet-lab procedures on single-cell RNA-Seq and provided technical support with bulk RNA sequencing. R.R.G. provided guidance and training on the *in vivo* experiments. D.G., S.R., A.S.-M., A.U.-O., M.U. and A.L.-I. generated the Basurto cohort and provided biological specimens. E.G.-B., A.R., J.T., D.J. and A.M. performed patient selection, tissue obtention and clinical annotation of the Morales Meseguer cohort. E.G.-B. supervised A.R., J.T., D.J. and A.M. N.M.-M., J.C.-M., L.B.-B., N.H., A.S. and H.V.S. performed the tissue preparation for single-cell experiments. J.M. recruited mHNPC patients in the single-cell validation cohort. M.G. supervised the single-cell preparation. A.C. conceived the study and supervised the execution of the project. I.M. and A.C. wrote the manuscript. All authors have read and approved the final version of the manuscript.

