## Supplementary_Figures for "The transcriptional landscape of metastatic hormone-naïve prostate cancer"

**Supplementary Figure and Table Legends**

**Supplementary Figure 1. A.** Principal component analysis of the transcriptomic data of the bulk discovery cohort. **B-E**. Heatmap and unsupervised clustering of top 50 differentially expressed genes (from the bulk discovery cohort) in four different primary tumour cohorts with available Gleason scores.

**Supplementary Figure 2. A.** Analysis of network topology for different soft-thresholding powers (WGCNA). The left panel shows the scale-free fit index (y-axis) as a function of the soft-thresholding power (x-axis). The right panel shows the mean connectivity (y-axis) as a function of the soft-thresholding power (x-axis). **B.** Clustering dendrogram of all genes based on the hierarchical clustering of adjacency-based dissimilarity. The coloured row below the dendrogram indicates module membership. **C.** Top: Hierarchical clustering dendrogram of module eigengenes (labeled by their colours). Bottom: Heatmap of the adjacencies in the eigengene network including the trait weight. Each row and column correspond to one module eigengene (labeled by colour) or weight. Blue represents low adjacency (negative correlation) and red represents high adjacency (positive correlation). **D.** WGCNA modules and their association with metastatic hormone-naïve (mHNPC) and localized (LPC) status and other biological and clinical variables. The percentage of DEGs in each module is also shown, as well as the p-value of the differences in intramodular connectivity between DEGs and non-DEGs per module (Wilcoxon test). **E.** Unsupervised hierarchical clustering of transcriptome-wide patterns of LPC and mHNPC prostate cancer primary tumours in the bulk validation dataset. **F.** Principal component analysis of the whole transcriptome in the bulk validation dataset. **G.** Volcano plot of the differential expression analysis in the validation cohort. Dots are coloured according to the WGCNA modules identified in the discovery cohort. **H.** Correlation between the $\log_2$ fold changes of the differential expression analyses in the discovery (y-axis) and validation (x-axis) cohorts for DEGs in both datasets. Each gene is coloured according to module membership in the discovery WGCNA analysis.

**Supplementary Figure 3. A**. Tumour purity and cell-type deconvolution inference of stromal and immune components in the discovery bulk dataset obtained from ESTIMATE. P-values from Wilcoxon test. **B.** Variance partition analyses of different biological and technical variables in the bulk discovery cohort. **C.** Tumour purity prediction and cell-type deconvolution results of stromal and immune components in the validation cohort. P-values from Wilcoxon test (two-tailed test for discovery and one-tailed for validation datasets, respectively). **D**. Variance partition analyses of different biological and technical variables in the bulk validation cohort.

**Supplementary Figure 4. A.** Dot plot displaying the expression levels of marker genes for cluster annotations in the discovery single-cell cohort. **B.** Inference of diploid (normal), aneuploid (tumour cells) or not defined based on COPYKAT in the discovery single-cell cohort. **C.** Cord diagram of the mHNPC-enriched ligands involved in cluster 6 and whole stroma (including immune stroma) communication that belong to purple module in the discovery single-cell cohort. Different stromal and immune clusters were merged into categories for visualization. **D.** Spearman correlation of the ligands identified in the single-cell validation dataset (cluster 6 with whole stroma) with transcription factors in mHNPC patients using the bulk discovery cohort. **E.** Spearman correlation of the ligands identified in the single-cell validation dataset (cluster 6 with whole stroma) with transcription factors in mHNPC patients using the bulk validation cohort. **F** Visualization of single-cell validation dataset (37,155 cells) using Uniform Manifold Approximation and Projection (UMAP). Colour codes for the assignment of each cell to different clusters by graph-based clustering (Leiden algorithm). **G.** Dot plot displaying the expression levels of marker genes for the clusters in the validation single-cell dataset. **H.** Inference of diploid (normal cells) and tumour cells (aneuploid) cells based on COPYKAT in the validation single-cell cohort. **I.** REACTOME pathway enrichment results for the 28 genes associated with cluster 6 in discovery dataset (i.e. markers that distinguish cluster 6 from other epithelial cells). **J.** Correlation analyses between the fold changes of differential gene expression (mHNPC vs LPC) in bulk and each single-cell cluster in the validation cohort for genes associated to the WGCNA purple module (minimum $|\log_2(\text{fold change})| > 0.6$). **K.** Cord diagram of mHNPC-enriched ligands involved in cluster 12-stromal and stroma (including immune) communication that belong to the purple module. Different stromal and immune clusters were merged into categories for visualization. **L.** Spearman correlation of the purple ligands identified in the single-cell validation dataset (cluster 12 with whole stroma) with transcription factors in mHNPC patients using bulk discovery cohort. **M.** Spearman correlation of the ligands identified in the single-cell validation dataset (cluster 12 with whole stroma) with transcription factors in mHNPC patients using the bulk validation cohort.

**Supplementary Figure 5. A.** Expression of two known SOX11 targets (PMID: 3396942) upon 3HA-SOX11 exogenous expression (n=3) in DU145 cells. One-sample t-test. **B.** Protein level changes upon 3HA-SOX11 exogenous expression in DU145 cells (n=4). Average $\text{Log}_2$ (ratio) of each protein is plotted against its $-\text{Log}_{10}$ (p-value). **C.** Left, dot plot of leg metastatic burden (day 51). One-tail, Mann Whitney test. Middle stacked bar plot of leg metastasis appearance. One-tail, Fisher test. Right, representative images of leg metastasis. **D**. Dot plot of the quantification of the ventral prostate burden (day 51). One-tail, Mann Whitney test. **E**. Relative cell growth upon 3HA-SOX11 exogenous expression in DU145 cells (n=3). One-sample t-test. **F.** KEGG pathway enrichment analyses of the genes upregulated upon *SOX11* overexpression (bulk RNA-Seq). **G.** Disease-free survival plots for SOX11 expression in different PC patient cohorts **H**. Disease-free survival plots for SOX11 activity signature in different non-mHNPC patient cohorts. **I.** Disease-free survival plots for the 13 ligands from the purple gene module involved in cancer cell-stroma crosstalk in different PC patient cohorts. p-values from long-rank test.

**Supplementary Table 1.** Patient characteristics of the four cohorts used in this study, including age, ISUP grading and metastasis classification (LPC: M0 and mHNPC: M1). The patients from the bulk cohorts were recruited in the Basurto University Hospital (Bilbao) and

Morales Meseguer Murcia in Spain (discovery and validation datasets, respectively). The patients for the single-cell validation dataset were obtained from Basurto University Hospital in Bilbao (LPC) and Vall d'Hebron Oncology Institute in Barcelona (mHNPC). See methods for more details.

**Supplementary Table 2.** Differential expression and WGCNA results on bulk discovery cohort.

**Supplementary Table 3.** Enrichment results of differentially expressed genes.

**Supplementary Table 4.** References and clinical characteristics of the different public prostate cancer datasets used in this study.

**Supplementary Table 5.** Results of the gene fusion analyses in the discovery bulk RNA-Seq cohort.

**Supplementary Table 6.** Association results of different WGCA modules with tumour type, age, RNA quality, PSA and Gleason grading.

**Supplementary Table 7**. Results of differential expression analyses of the bulk validation cohort. The table also contains normalized counts used as input in those analyses.

**Supplementary Table 8**. Results of ESTIMATE for discovery and validation bulk RNA-Seq datasets.

**Supplementary Table 9.** Differential expression results (pseudo-bulk) for single-cell discovery dataset per cluster.

**Supplementary Table 10.** Correlation analyses (Spearman's) between the fold changes (mHNPC vs LPC, minimum $|\log_2(\text{fold change})| >0.6$)) in bulk and single-cell for the seven modules associated with mHNPC according to WGCNA analysis. Darkmagenta did not show any gene passing the minimum fold change criteria. Results for all single-cell clusters are shown.

**Supplementary Table 11.** Results of cancer cell-stromal communication in single-cell datasets using NicheNet. In the discovery single-cell cohort the sender cluster was cluster 6 and stromal cells were the receivers. In the validation dataset the sender cluster was cluster

12 and the receivers were the stromal cells. Results provided for LPC and mHNPC niches per cohort. Results for both whole stroma and non-immune stroma are provided.

**Supplementary Table 12.** Results for the correlation between purple ligands with transcription factors in bulk transcriptome data from mHNPC patients from the discovery and validation bulk cohorts.

**Supplementary Table 13.** Differential protein abundance results of SOX11 overexpressing DU145 cells vs control cells.

**Supplementary Table 14.** Differential expression results of SOX11 overexpressing DU145 cells vs control cells and functional enrichment results for the differentially expressed genes.

**Supplementary Figure 1**

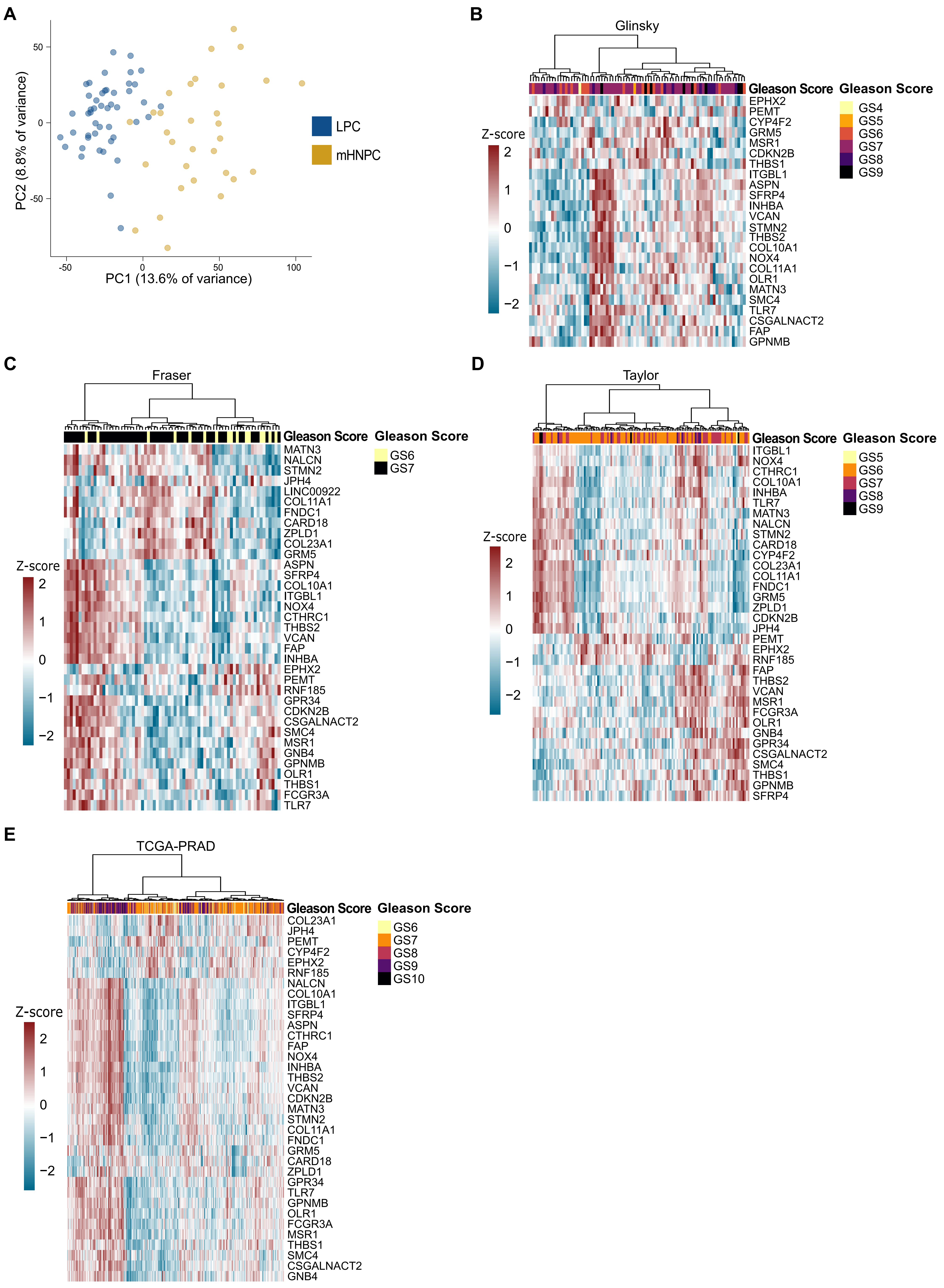

**Supplementary Figure 2**

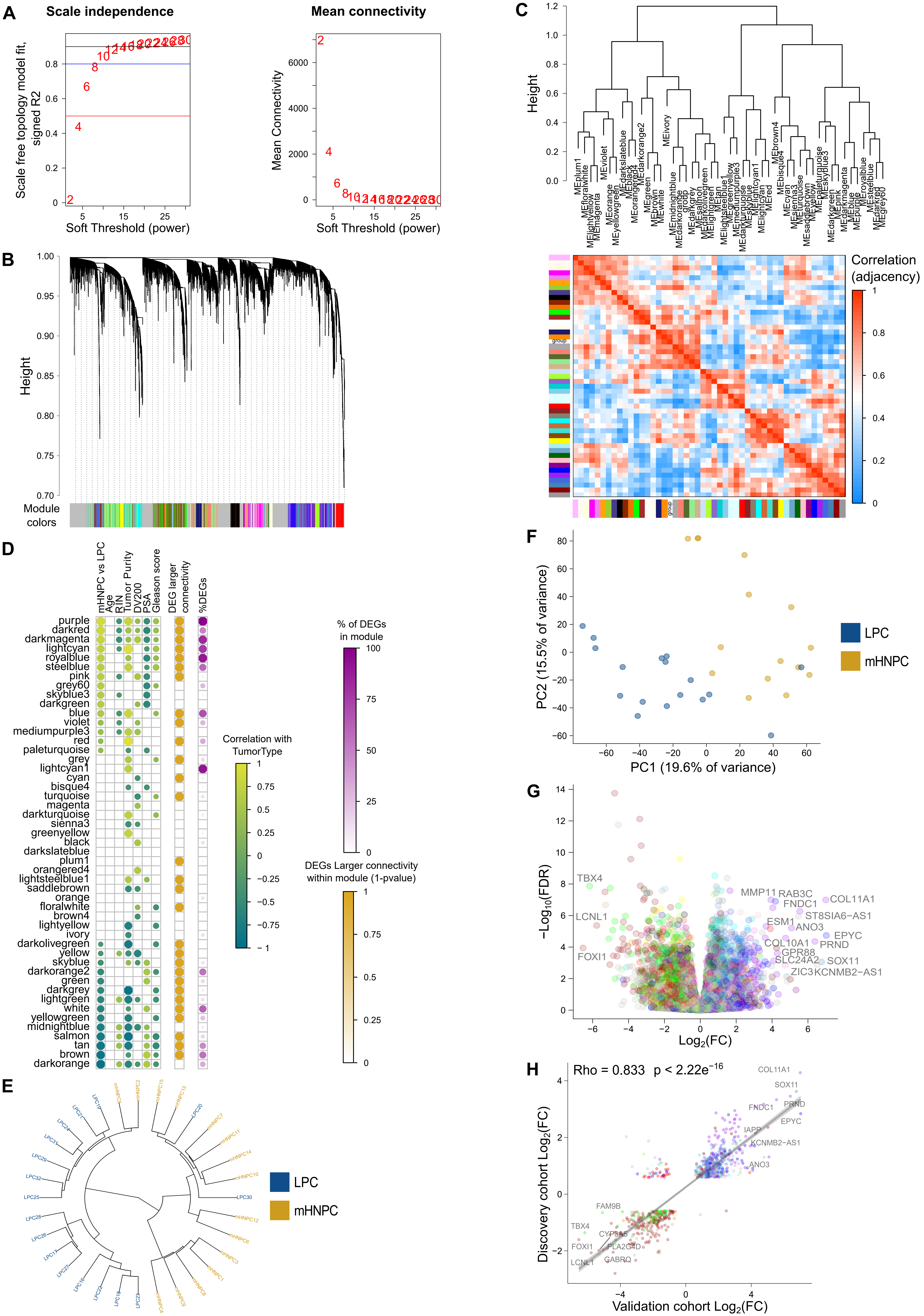

### Supplementary Figure 3

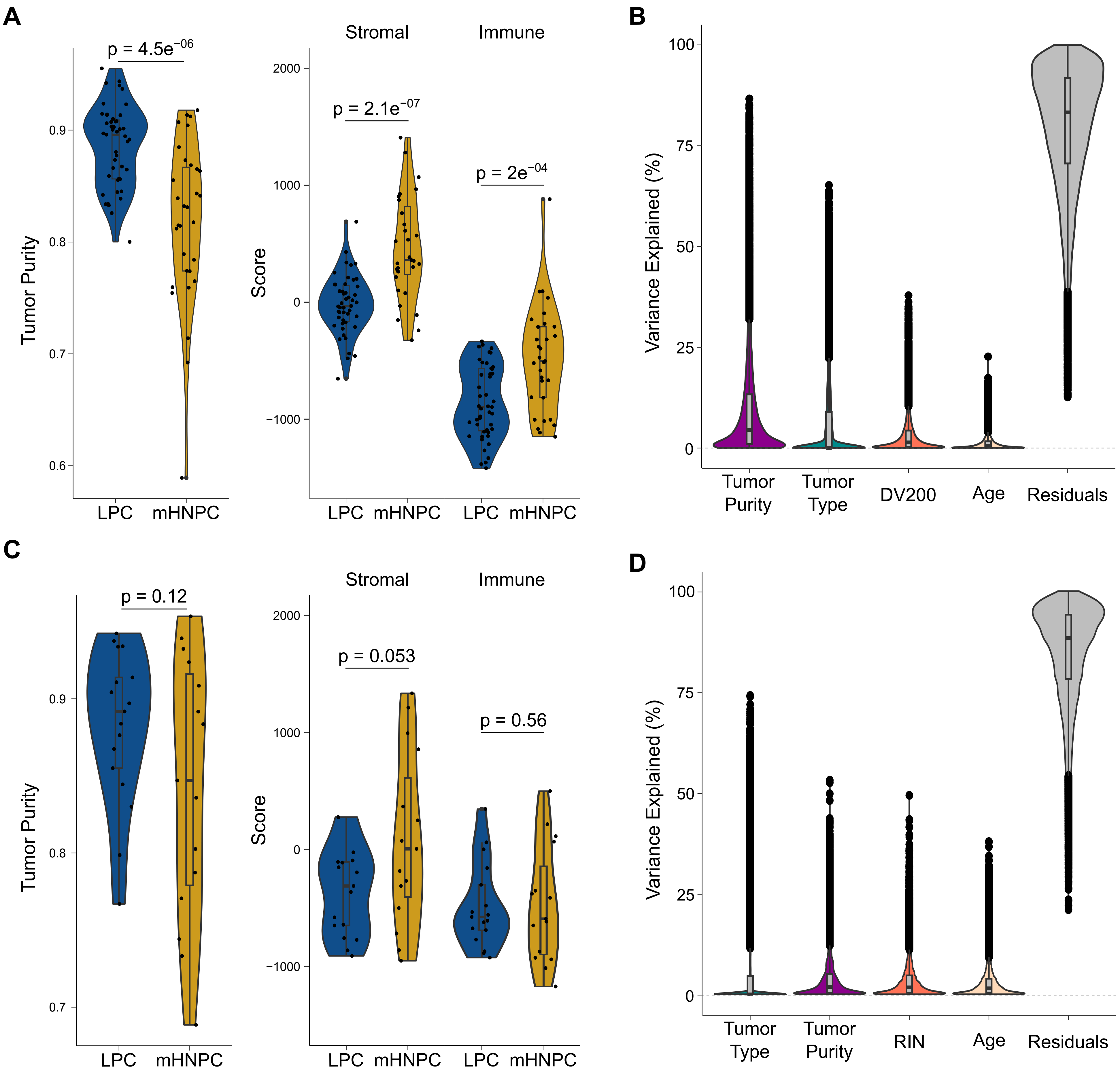

### Supplementary Figure 4

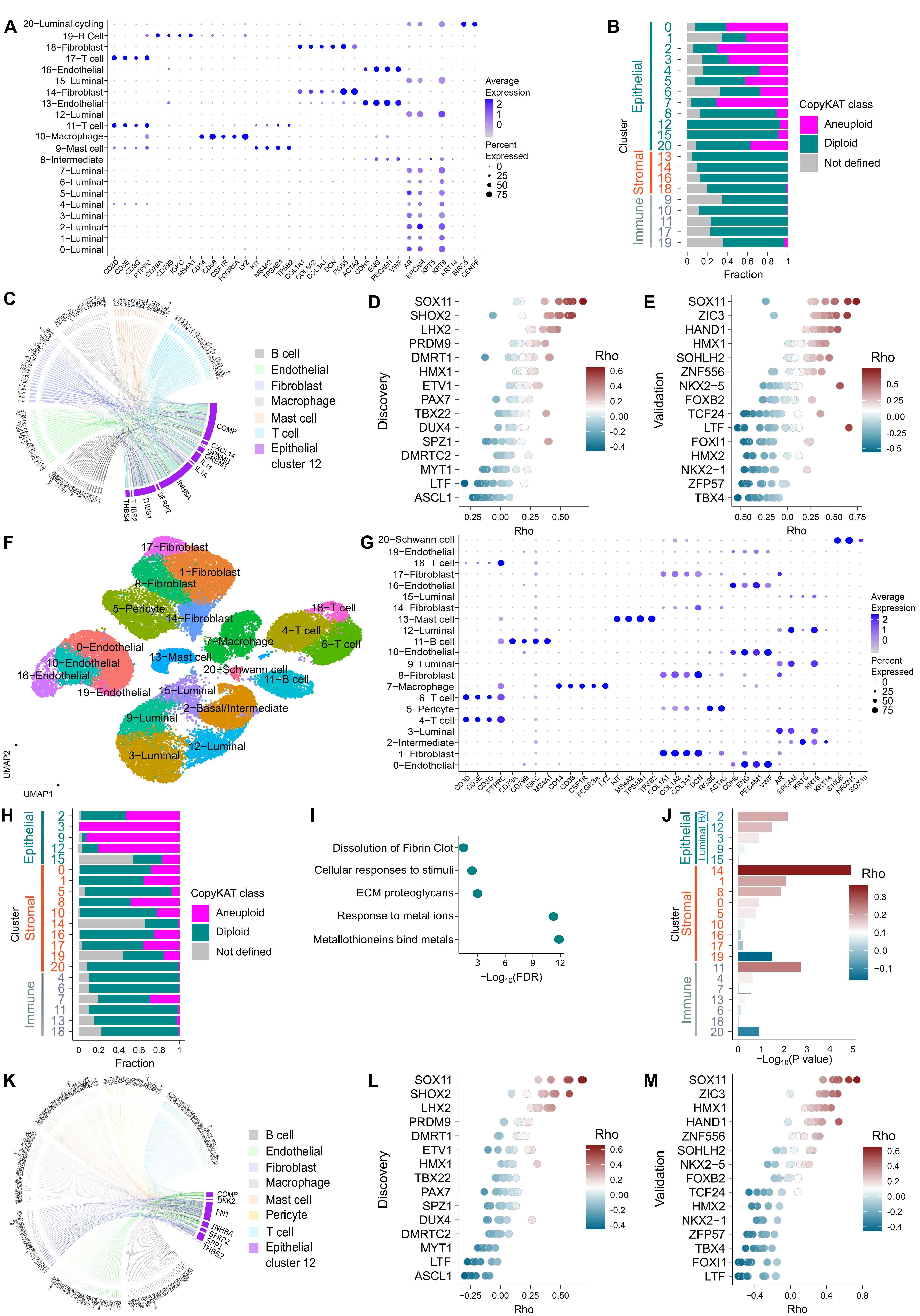

#### Supplementary Figure 5

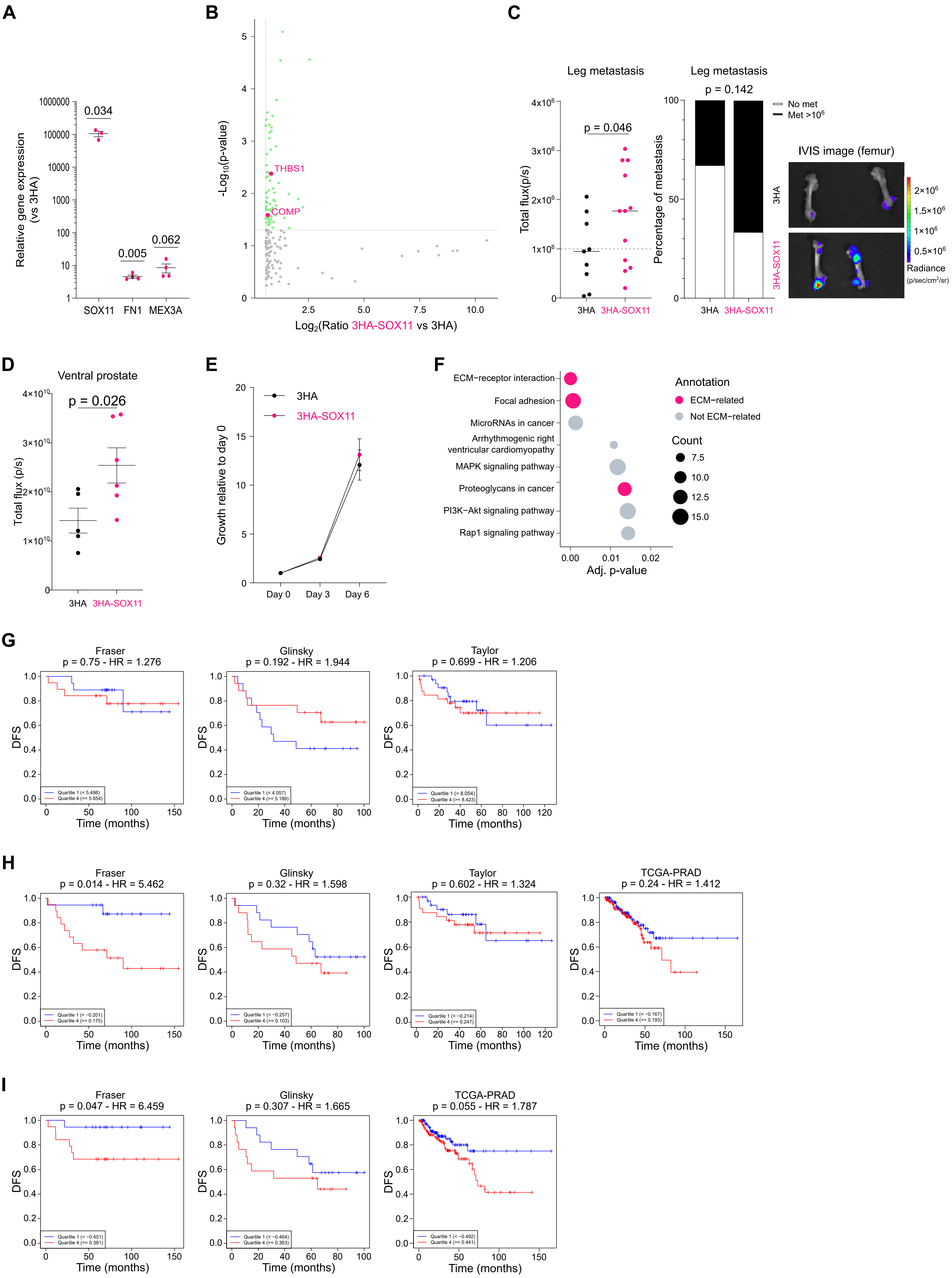
